# A survey of Chinese pig farms and human healthcare isolates reveals separate human and animal MRSA populations

**DOI:** 10.1101/2021.08.02.454852

**Authors:** Geng Zou, Marta Matuszewska, Ming Jia, Jianwei Zhou, Xiaoliang Ba, Juan Duan, Caishi Zhang, Jian Zhao, Meng Tao, Jingyan Fan, Xiangming Zhang, Wenping Jin, Tianpen Cui, Xianyu Zeng, Min Jia, Xiaojuan Qian, Chao Huang, Wenxiao Zhuo, Zhiming Yao, Lijun Zhang, Shaowen Li, Lu Li, Qi Huang, Bin Wu, Huanchun Chen, Alexander W. Tucker, Andrew J. Grant, Mark A. Holmes, Rui Zhou

## Abstract

There has been increasing concern that the overuse of antibiotics in livestock farming is contributing to the burden of antimicrobial resistance in people. Farmed animals in Europe and North America, particularly pigs, provide a reservoir for livestock-associated methicillin-resistant *Staphylococcus aureus* (LA-MRSA, ST398) found in people. This study was designed to investigate the contribution of MRSA from Chinese pig farms to human infection and carriage.A collection of 603 *S. aureus* were isolated from 55 pig farms and 4 hospitals (MRSA= 285, 198; MSSA= 50, 70) in central China, a high pig farming density area, during 2017-2018. CC9 MRSA accounting for 93% of all farm MRSA isolates, while no was found in hospitals. ST398 isolates were found on three farms (n = 23) and three hospitals (n = 12). None of the ST398 from this study belong to the livestock clade of the LA-MRSA commonly found in Europe and North America. The hospital ST398 MRSA isolates formed a clade that was clearly separate from the farm ST398 MRSA and MSSA isolates, and all possessed human immune evasion cluster genes which were absent from all the pig farm ST398 isolates. Despite the presence of high levels of MRSA found on Chinese pig farms we found no evidence of them spilling over to the human population. Nevertheless, the ST398 MRSA obtained from human samples appear to be part of a widely distributed lineage in China. And the new animal adapted ST398 lineage that emerged in China should also be alarmed.

**Importance:** We disclosed the fact that although the high MRSA positive rate in Chinese hospitals and pig farms should be alarmed, they might be two separate issues. The new CC398 clades we identified highlight that the host adaption of the MRSA lineage is kept changing. These results suggest that continued surveillance of MRSA in livestock is necessary. We found that the pig farm MRSA isolates had unique antimicrobial resistance genes while most of the hospital MRSA isolates had human immune evasion cluster genes. These features could be used to distinguish the pig farm associated S. aureus in clinical laboratories. The policies of reducing antimicrobials use in livestock were implemented in China since 2020. Our study described the situation of MRSA populations in pig farms and hospitals in Central China before 2020, which provides a potential opportunity for future studies to evaluate the effects of the policies.

## Introduction

*Staphylococcus aureus* (*S. aureus*) is an opportunist human pathogen that can also colonize and infect many species of animals and adapt to diverse environmental conditions ^1^. Methicillin-resistant *S. aureus* (MRSA) are *S. aureus* lineages that have acquired the *mecA* or *mecC* genes through horizontal gene transfer which makes them highly resistant to nearly all β-lactam antibiotics ^2^.

The zoonotic potential of MRSA in livestock and wild animals has been widely reported ^3^. An early report of livestock associated MRSA (LA-MRSA) described an infection in a baby living on a farm where the pigs were infected with MRSA Sequence Type (ST)398, in the Netherlands in 2005 ^4^. This LA-MRSA lineage is now widely distributed in pigs and other farm animal populations in Europe and North America ^5,6^. A recent national report of AMR in Denmark gave the prevalence of LA-MRSA in finishing pigs in randomly selected herds as 88% with ST398 accounting for 16% of all human MRSA infections which raises considerable concerns about antibiotic stewardship in agriculture and the threat to public health^7^.

Phylogenetic studies have shown that LA-MRSA ST398 originated from a human MSSA lineage and jumped to livestock, acquiring tetracycline and methicillin resistance ^8,9^. These two clades can be distinguished on the basis of canonical single-nucleotide polymorphisms (SNPs) ^10^.

The LA-MRSA clonal lineage most frequently reported in Asian livestock is ST9 with closely related isolates from people and livestock reported in the literature ^11, 12^. This ST is also common in European and North American pig farms almost exclusively as a MSSA. The origins of this lineage are unclear although it is common in historic collections of livestock isolates suggesting an animal origin ^13, 14^.

In addition to ST398 and ST9 MRSA, among other lineages (CC1, CC5, CC8, CC59, CC97, CC130 and CC425) have been found in livestock with evidence that they are also involved in zoonotic or anthroponotic transmission (host switching) ^15,16^.

Previous studies looking at LA-MRSA in China have shown that ST9 is the dominant lineage in pigs with ST398 MRSA being less commonly detected ^17,18,19^. Studies of human isolates have identified ST398 MRSA as a cause of significant disease in patients from Chinese hospitals^20^. Genomic studies of human isolates have indicated that some of these isolates are from the human ST398 lineage having acquired a *mecA* gene in a distinctive SCC*mec* V variant ^21^.

In order to assess the contribution of pig farms to the human burden of MRSA, a study was designed to enable a genomic study of MRSA isolates from human and animal hosts. Isolates of *S. aureus* were collected contemporaneously from pig farms and hospitals in the same region in China and subjected to DNA sequencing. A phylogenetic analysis was performed in order to establish if the human isolates were likely to be of livestock origin.

## Results

### The MRSA populations of pig farms and hospitals consisted of different clonal lineages

In the samples collected from 55 pig farms (pig nasal swabs = 2416, farm worker nasal swabs = 361, pig farm dust samples = 291), 335 *S. aureus* isolates were identified and sequenced (based on phenotypic antibiotic susceptibility 285 of them were MRSA). MRSA were isolated from 34/55 pig farms (62%). CC9 MRSA were present in all positive farms where more than one MRSA isolate was obtained and accounted for 97% (277/285) of all farm MRSA isolates. ST398 isolates were found on 3 farms: one ST398 MRSA was isolated from a farm worker on one farm, two ST398 MRSA isolates were obtained from a pig from another farm, and a number of ST398 MSSA were isolated from pigs, environmental samples and a farm worker on the third farm. The remaining 5 farm MRSA isolates comprised one ST1 (from a pig sample) and 4 ST59 (1 from a pig and 3 from farm worker swabs) (Figure S1, Table S1). 50 MSSA pig farm isolates included 16 ST9, 5 ST1, 4 ST5, 20 ST398, and 5 ST128. (Table 1).

**Table 1.**
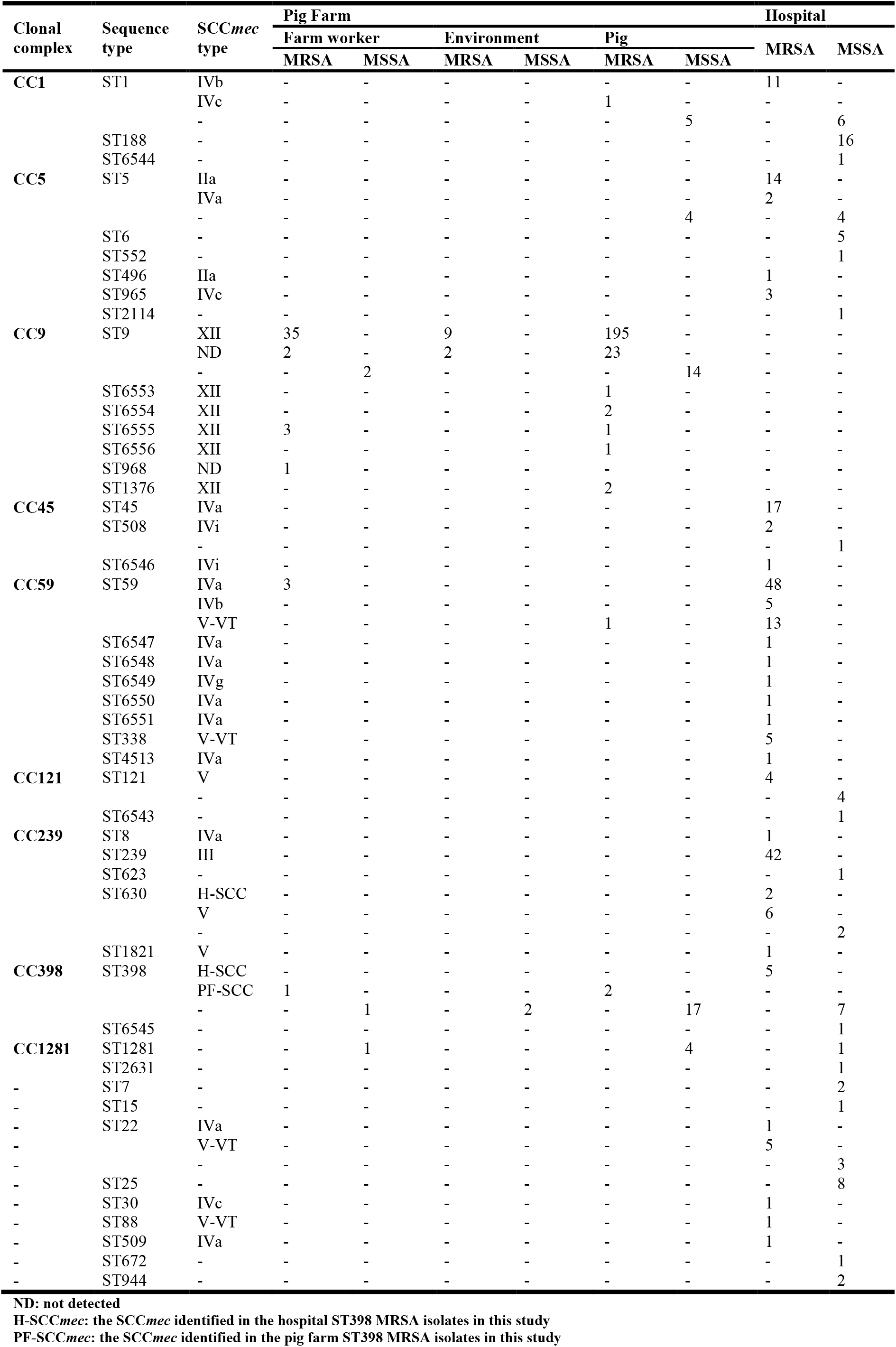
The distribution of the whole *S. aureus* collection according to different sample sources.

A total of 268 *S. aureus* isolates were collected from four hospitals (MRSA = 198, MSSA = 70) and 39 different STs were identified. The three most common STs in the hospital MRSA populations were ST59 (66/198), ST239 (42/198) and ST45 (17/198). Five ST398 MRSA isolates were identified from three hospitals, including two city hospitals and a county hospital, seven ST398 MSSA isolates were also identified in the county hospital (Table 1). Notably there were no ST9 isolates (MRSA or MSSA) found in the Chinese hospitals.

### Phylogenetic analysis identifies separate human and livestock ST398 clades in China

Phylogenetic studies can provide evidence about the population structure and rates of transmission between different hosts. Where genomes from different hosts are mixed in each clade, this indicates jumps between host species. Where strains from different hosts form distinct clades, this suggests that distinct populations are being maintained in different hosts, and that host jumps are rare^16^. A phylogenetic analysis was performed to examine the population structure of ST398. A total of 115 Chinese CC398 isolates, including 36 isolates from this study, 37 isolates collected from NCBI genome database and 42 isolates from hospital patients collected in a previous study ^21^ were used to create an alignment against the reference genome 0213-M-4A (a finished pig ST398 MRSA genome generated in this study) from which a maximum likelihood tree was constructed (Figure 1).

**Figure 1.**
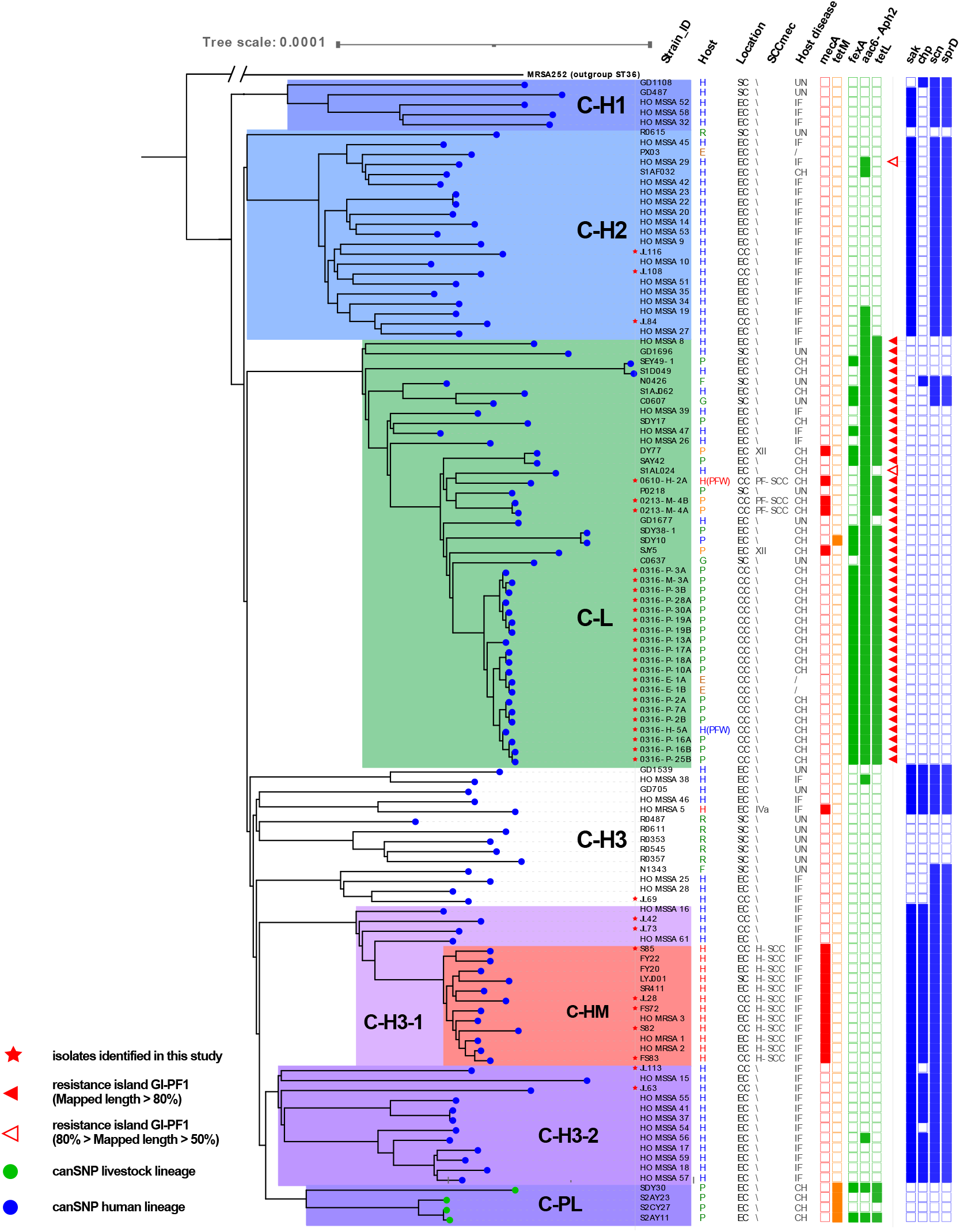
The phylogenetic tree illustrates the relationship of the Chinese CC398 isolates. Based on whole genome SNPs, a phylogenetic tree including 115 Chinese CC398 isolates (107 ST398, 8 other STs) from this study and other published studies is shown. An MRSA ST36 strain (GenBank: BX571856.1) was used as the out group. Location: EC, East China; SC, South China; CC, Central China. Host: B, Bovine; H, Human; P, Porcine; F, Feline; R, Rodent; G, Galline/Chicken; E, Environment; H(PFW), Human from a pig farm worker sample. SCC*mec*: For each isolate, the SCC*mec* type was determined if more than 70% of SCC*mec* elements could be detected in the genome. Some *mecA* negative isolates were found to have SCC*mec* elements using this method. Host disease: UN, Unknown; IF, Infected with *S. aureus*; CH, Clinically healthy. ARGs and VFs: the filled squares indicate the presence and empty squares the absence of individual genes indicated at the head of each column.

With the exception of 4 MSSA from pigs in eastern China (Clade C-PL), all the Chinese CC398 MRSA and MSSA isolates (from both livestock and humans) were shown to fall within the diversity of the previously identified ‘human’ lineage as defined by the published canonical SNPs (Figure 1). The Clade C-PL isolates are contained within the previously identified ‘livestock’ lineage and carry a *tet*M gene which is associated with the ‘livestock’ lineage^8^. All the human hospital MRSA ST398 isolates from this study formed a discrete cluster with other MRSA in the collection from a broad geographical range in China (Clade C-HM). No pig farm (or other animal) isolates contributed to this clade and they all possessed a full set of the human immune evasion cluster genes. The pig farm ST398 MRSA isolates fell within the diversity of another separate clade (Clade C-L) which was comprised of the majority of animal MSSA isolates together with some human MSSA isolates and 5 MRSA (1 farm worker from this study 4 from pigs). All isolates in Clade C-L lacked a full set of the human immune evasion cluster genes and contained a characteristic set of antimicrobial resistance genes (see below). The pig farm MSSA CC398 isolates identified in this study (which all came from a single farm) clustered within Clade C-L. The 8 hospital MSSA CC398 isolates from this study belonged to 2 separate clades (Clade C-H2 and C-H3). Three isolates (JL116, JL108 and JL84) were in the Clade C-H2 separated from most of the other human MSSA. Two (JL42 and JL73) of the other 5 isolates fell in the Clade C-H3-1 containing human hospital MRSA (Clade C-HM) from this study suggesting that SCC*mec* had been acquired by this human-adapted lineage. Two (JL113 and JL63) of the remaining 3 isolates belonged to the Clade C-H3-2 consisting entirely of human MSSA which had a relatively close relationship to the ‘livestock’ lineage (Clade C-PL).

A time-measured phylogenetic tree using an international collection of ST398 isolates was created using BEAST1.8.2 ^22^. After removing all isolates without date, host, and location information in the selection process (see methods), a total of 207 ST398 isolates (11 from this study, and 196 from the NCBI genome collection) were used for the analysis. The collection included isolates collected from around the world over the last 15 years. A time annotated tree is shown in Figure 2. The tree shows an ancestral clade at the base of the tree consisting of mainly human MSSA ST398 isolates from Europe and North America, which also includes a Chinese clade (Clade C-H1). The well documented prevalent LA-MRSA clade (the ‘livestock’ clade; possessing the appropriate canonical SNPs) appears to be the most recently emerged clade. It includes just 2 Chinese isolates (from the Clade C-LP in Figure1) from pigs and is defined by a most recent common ancestor (MRCA) dating back to 1964 (95% highest posterior density interval [HPD] = [1956, 1972]). With a MRCA estimate of 1940 (95%HPD = [1926, 1954]) the vast majority of Chinese isolates and the ‘livestock’ clade emerged from the ancestral ST398 clade. Three Chinese clades (Clade CH2, C-L and C-H3) all appear to separate at the same time with the MRCA dated at 1950 (95%HPD = [1938, 1961]). Clade C-H2 consists mainly of human MSSA, clade C-L consists mainly of livestock MSSA (but also contains 2 MRSA from farms in this study), and clade C-H3 consists of another clade of human MSSA (Clade C-H3-1) from which emerges with the MRCA estimated at 2002, (95%HPD = [1999, 2005]) the clade of human hospital MRSA (Clade C-HM). An additional human MSSA Chinese clade (Clade C-H3-2) shares a MRCA with the ‘livestock’ clade dated at 1956 (95%HPD = [1946, 1966]).

**Figure 2.**
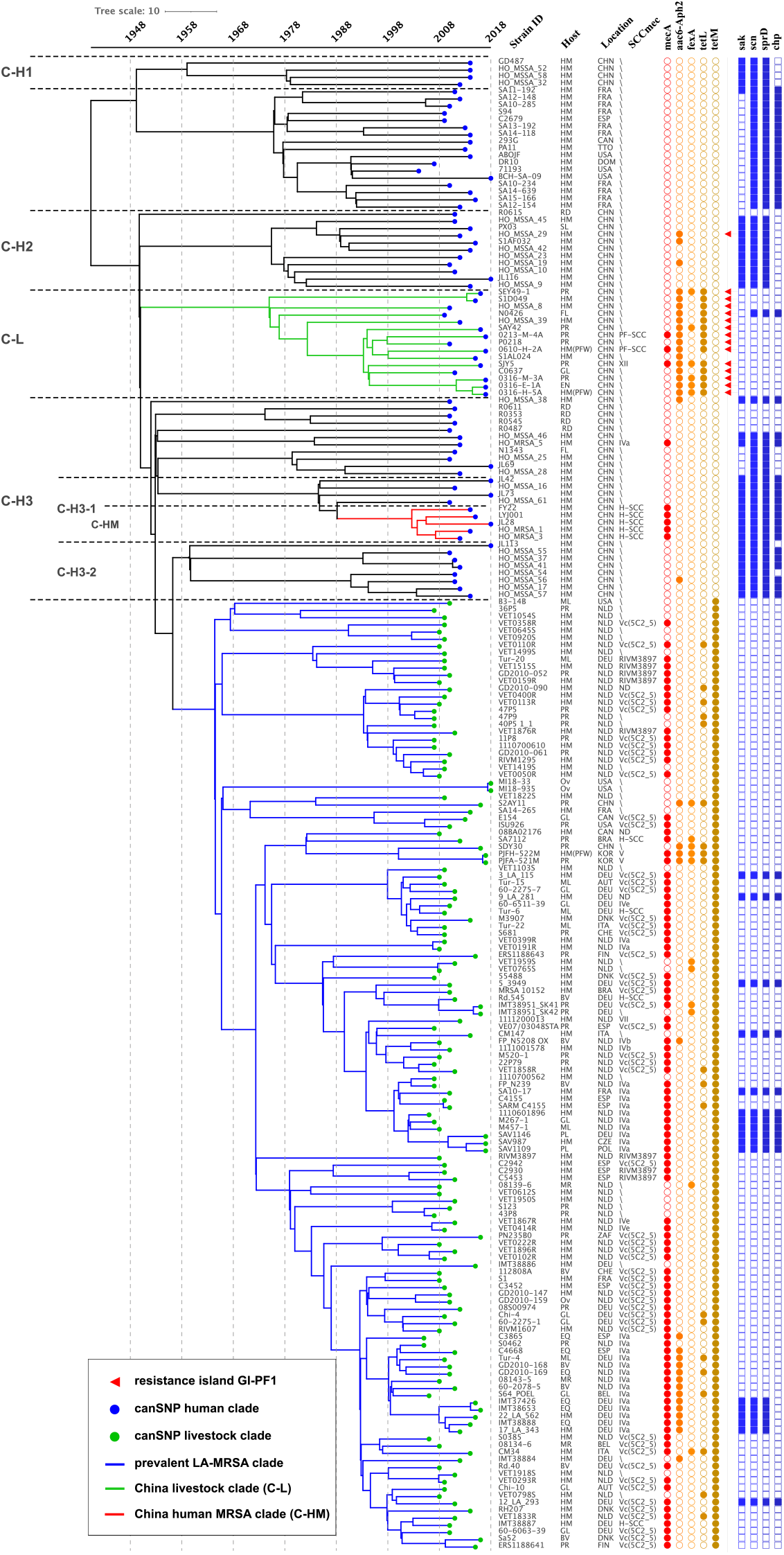
Phylogeny and molecular dating of the international ST398 isolates. A time-measured phylogenetic tree using an international collection of ST398 isolates was obtained using BEAST2. A total of 207 ST398 isolates (11 from this study, and 196 from the NCBI genome collection) were used in the analysis. Host: HM, Human; PR, Porcine; EN, Environment; HM(PFW), Human from a pig farm worker; MR, Murine; BV, Bovine; GL, Galline; ML, Meleagrine; EQ, Equine; OV, Ovine; FL, Feline; RD, Rodentia; PL, Poultry. Location: DNK, Denmark; USA, United States of America; NLD, Netherlands; CHN, China; BEL, Belgium; CAN, Canada; DEU, Germany; CHE, Switzerland; ESP, Spain; AUT, Austria; ITA, Italy; DMA, Dominica; BRA, Brazil; TTO, Trinidad & Tobago; FIN, Finland; KOR, South Korea; ZAF, South Africa; FRA, France; POL, Poland; CZE, Czech Republic. SCC*mec*: For each isolate, the SCC*mec* type was determined if more than 70% of SCC*mec* elements could be detected in the genome.

### The pig farm CC9 MRSA may be of importance to public health

The phylogenetic tree of CC9 *S. aureus* from this study and previous studies roughly separates into three CC9 populations (Figure 3). Towards the root of the tree there is a clade of genomes that possess human immune evasion cluster (IEC) genes (*chp, scn* and *sak*) that includes two human clinical MRSA isolates from Chinese Taiwan with a SCC*mec* V variant. The vast majority of Chinese isolates form a “China pig-farm” clade, are mostly of animal origin, and lack the IEC genes. The China pig-farm clade shares a MRCA with a North American/European clade (the third population which includes an MRSA lineage with a type IVb SCC*mec*. The China pig-farm clade isolates are almost all MRSA with SCC*mec* XII and this clade contains all isolates from this study (145 included in this tree). In this clade, of the 45 isolates from other studies, nine were from human clinical cases. As the absence of human IEC genes and the possession of the farm associated antimicrobial resistance genes (ARGs) (*tetL, fexA* and *aac6-Aph2*) is a consistent feature in this clade it seems likely that the human clinical isolates probably originated from livestock as it is clear this lineage is found in human carriage samples including farm workers. Notably, there was a single isolate with the human IEC genes isolated from a bacteraemia in a patient from a hospital in Hangzhou (East China)^23^. Within the phylogeny this isolate is surrounded by pig farm isolates from this study.

**Figure 3.**
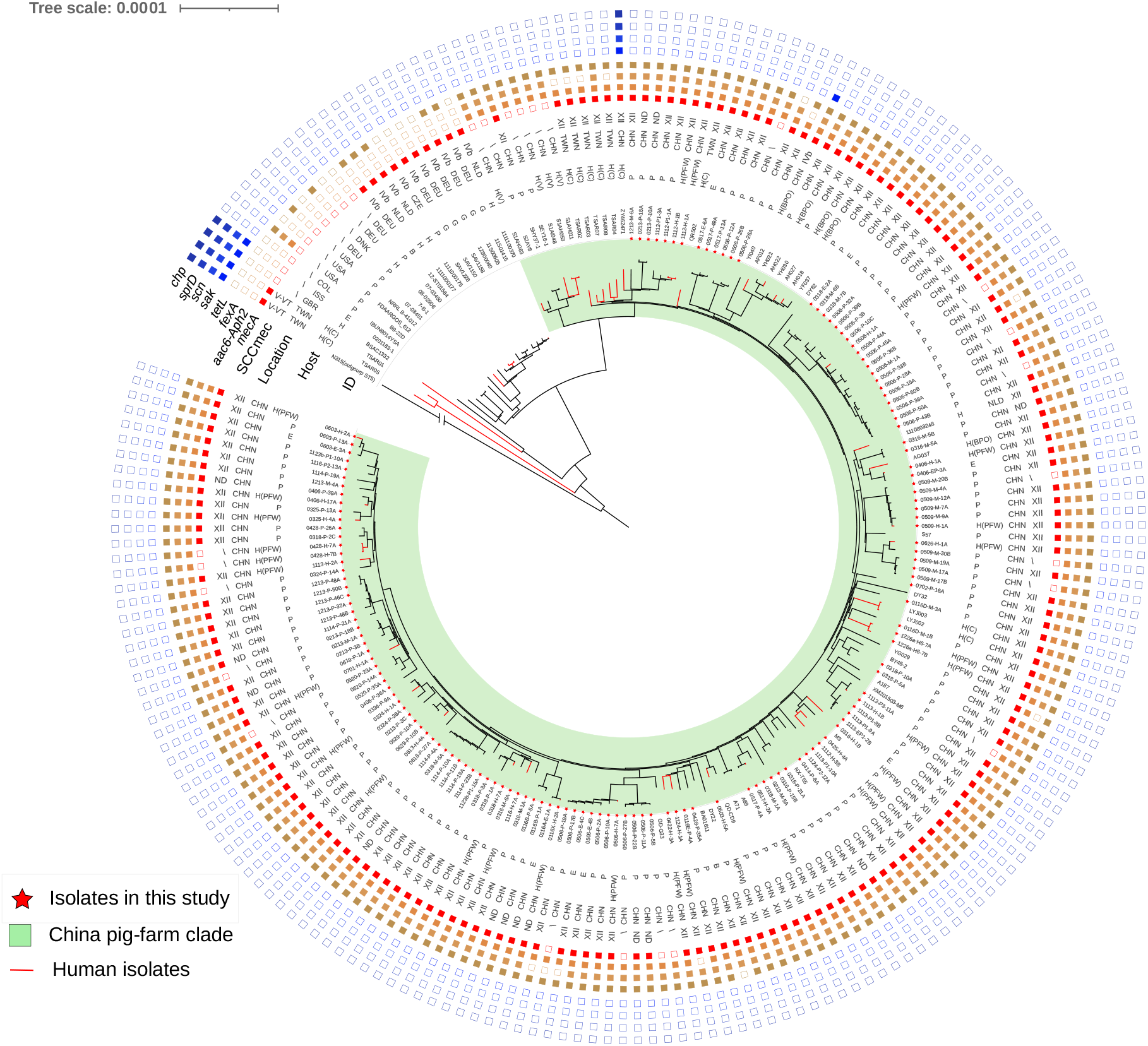
The phylogenetic tree illustrates the relationship of the international ST9 isolates. Based on whole genome SNPs, a phylogenetic tree consisting of 144 isolates selected from this study and 63 isolates from other reports is shown. A MRSA ST5 strain (GenBank: NC_002745.2) was used as the out group. Location: CHN, China; COL, Colombia; CZE, Czech Republic; DEU, Germany; DNK, Denmark; GBR, United Kingdom; ISS, International Space Station Air (U.S. Lab); NLD, Netherlands; TWN, Chinese Taiwan; USA, United States. Host: B, Bovine; E, Environment; G, Galline/ Poultry; H, Human; H(C), from a human *S. aureus* infection clinical sample; H(BPO), from a backyard pig farm owner sample; H(PFW), from a pig farm worker sample; H(V), from a village community sample. P, Porcine. SCC*mec*: ND, *mecA* positive but not detected published SCC*mec*. ARGs and VFs: the filled squares indicate the presence and empty squares the absence of individual genes indicated at the head of each column.

### Different patterns of antimicrobial resistance and virulence genes from pig farms and hospitals

An examination of ARGs and selected virulence factors (VFs) in *S. aureus* sequence data showed that 3 ARGs were frequently found in pig farm isolates (PFA-ARGs), *aac6-Aph2* (aminoglycosides ARG)^24^, *fexA* (phenicols ARG)^25^ and *tetL* (tetracyclines ARG)^26^; and 3 VFs frequently found in hospital isolates (HA-VFs), *scn* (*Staphylococcal* complement inhibitor)^27^, *sak* (staphylokinase)^27^ and *sprD* (small pathogenicity island RNAs)^28^ (Figure S2). The PFA-ARG *aac6-Aph2* was present in 9/10 clonal lineages of the pig farm MRSA population (98% of pig farm MRSA isolates). It was also found in 5/24 clonal lineages from hospital MRSA isolates particularly in ST239 (41/42) and ST5 (11/16). The ARGs, *fexA* and *tetL*, were not present in hospital MRSA isolates (Figure 4). The HA-VF *sak* was found in 76% (150/198) of hospital MRSA isolates accounting for 20/24 clonal lineages while 82% (162/198) harboured *scn* and 83% (164/198) harboured *sprD*. The HA-VFs were only present in ST1 and ST59 pig farm MRSA isolates, mostly from farm worker samples. In comparison to ST398 MRSA isolates from patients, ST398 isolates from pig and farm worker samples possessed ARGs *aac6-Aph2* and *tetL* but lacked HA-VFs. The single pig ST1 MRSA isolate in the collection harboured PFA-ARGs *aac6-Aph2* which was not present in the hospital ST1 MRSA isolates. Except that a couple of (2/66) hospital ST59 MRSA isolates carried the *aac6-Aph2*, no other PFA-ARG was detected in ST59 MRSA isolates. HA-VF *sak* was not detected in pig ST59 MRSA isolate nor in 11/66 hospital ST59 MRSA isolates (Figure 4).

**Figure 4.**
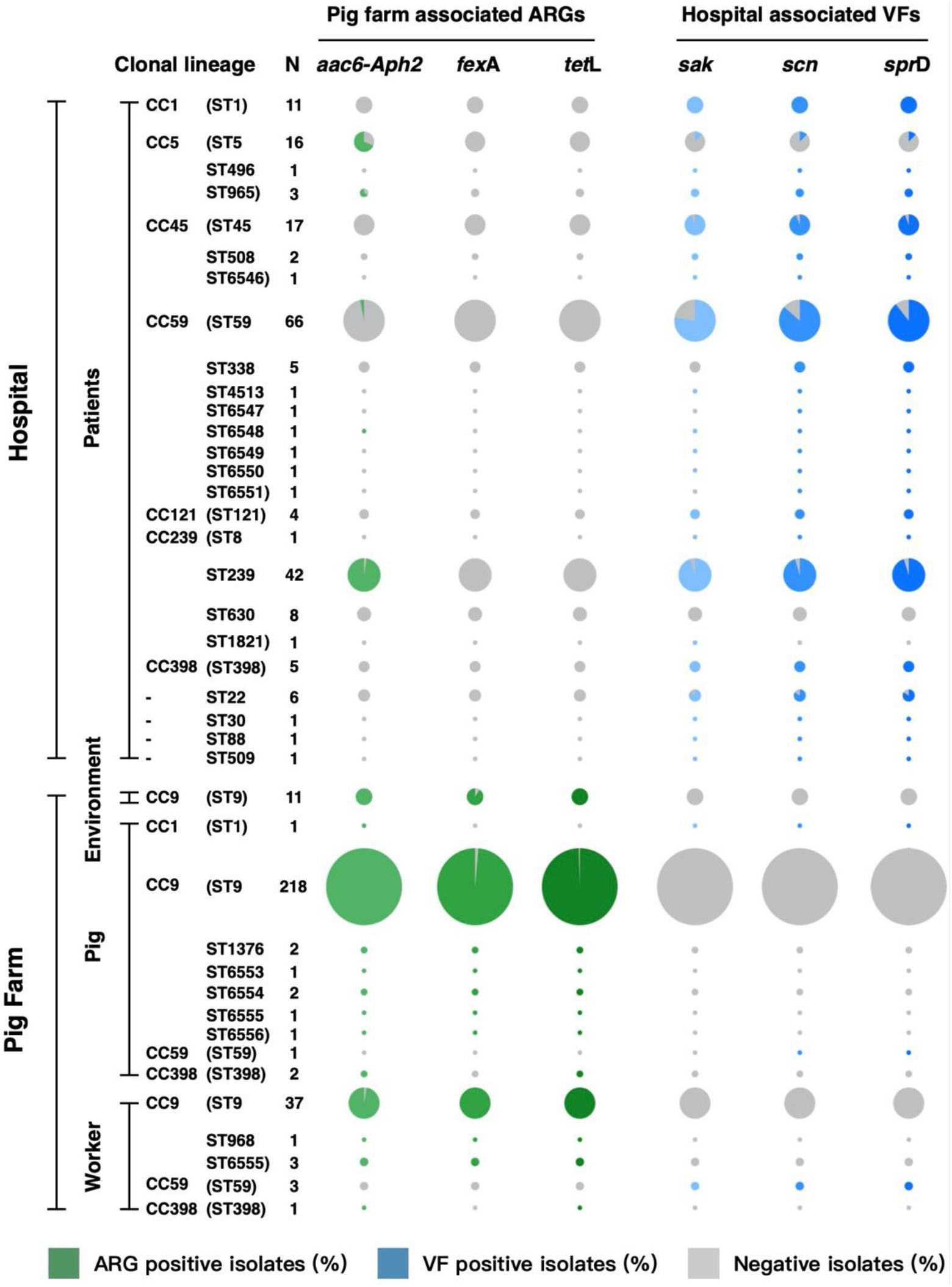
Distribution of pig farm associated antimicrobial resistance genes and hospital associated virulence factor genes among clonal lineages. The distribution of three antimicrobial resistance genes (ARGs) and three hospital associated virulence factor genes (VFs) among clonal lineages of all isolates is shown. The pie charts in each row, indicate the proportion of the isolates from each ST harboring individual ARGs or VFs as defined by the column, with the size of each pie chart proportional to the number of isolates.

### Mobile genetic elements harboured by MRSA ST398 isolates from pig farms and hospitals

An examination of the genomic context of the PFA-ARGs and HA-VFs is illustrated in Figure 5A. A transposon, Tn558, harbouring *fexA* was detected in the pig farm MSSA ST398 isolate, but not in the pig farm MRSA ST398 isolates or the hospital ST398 isolates. Tn558 could be found in all (313/313) of the *fexA* positive isolates isolated for this study.

**Figure 5.**
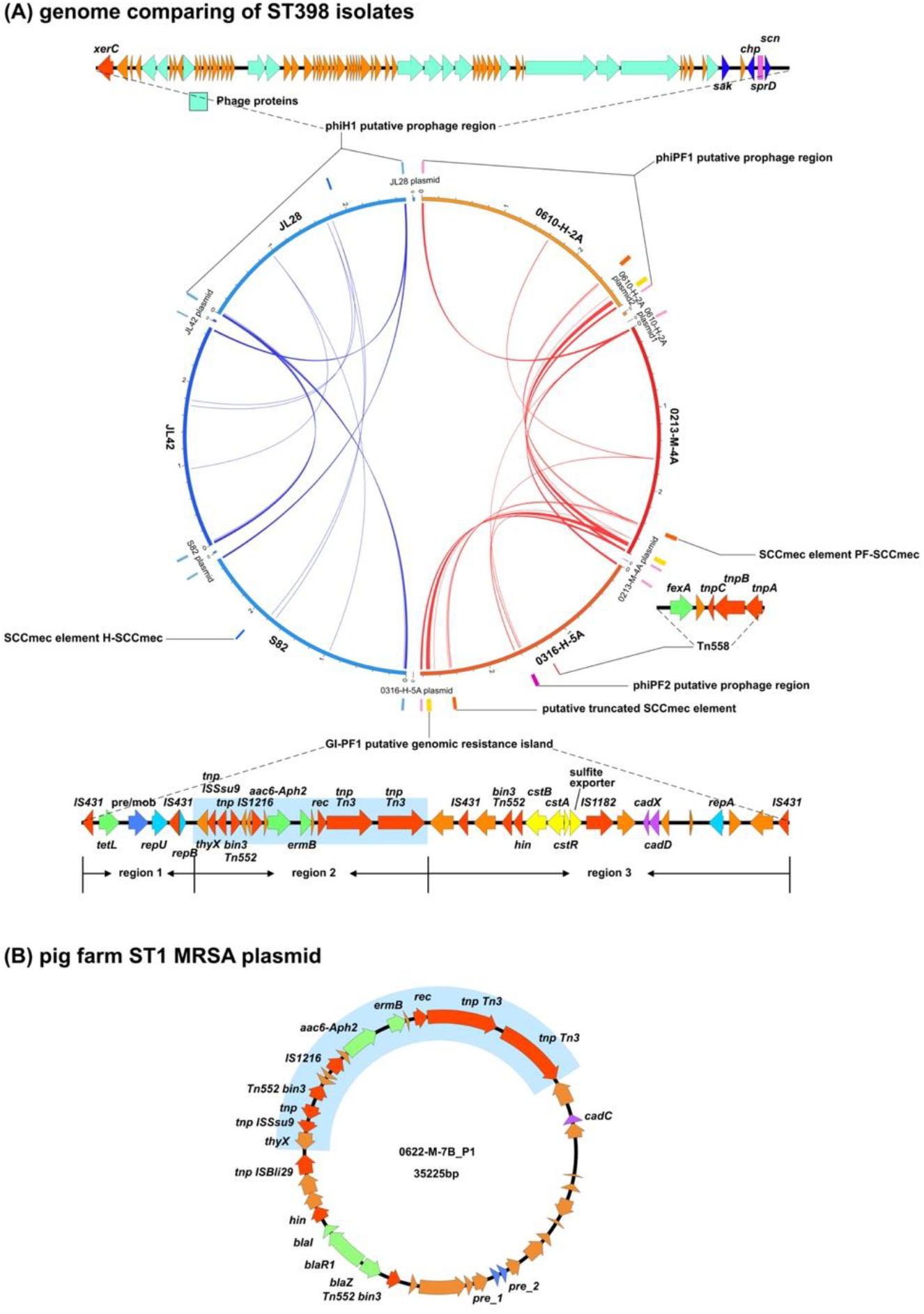
Genomic elements carrying pig farm associated antimicrobial resistance genes and hospital associated virulence factor genes. The genomic elements harboring the pig farm associated ARGs or hospital associated VFs were identified by comparing the ST398 isolates of pig farms and hospitals. (A) The distribution of the genomic elements in six ST398 completed genomes are shown: JL28, hospital ST398 MSSA isolate; JL42 and S82, hospital ST398 MRSA isolates; 0213-M-4A: pig farm MRSA isolate from a sow nasal sample; 0610-H-2A: pig farm MRSA isolate from a farm worker nasal sample; 0316-H-5A: pig farm MSSA isolate from a farm worker nasal sample. The genomic sequences shared only between the hospital isolates are indicated using blue curves, and genomic sequences shared only between the pig farm isolates are indicated using red curved lines. Diagrammatic representations of the genomic elements are illustrated, and their locations are shown adjacent to each genome included in the central circular figure. Individual transposon associated genes are shaded red, ARGs are shaded green and purple is used to show genes conferring resistance to metals. (B) The panel contains a diagram of a plasmid from the ST1 pig farm isolate 0622-M-7B. When compared to the ST1 MRSA isolates from the hospitals, the pig farm MRSA ST1 isolates had two additional ARGs, *aac6-Aph2* and *ermB*. These two ARGs were found in a plasmid with a sequence that was identical with region 2 of the GI-PF1 identified in the pig farm ST398 isolates (highlighted in blue).

A putative genetic resistance island, GI-PF1, carrying the ARGs *tetL* and *aac6-Aph2* as well as the erythromycin resistance gene *ermB*, appeared to be present in all the pig farm ST398 isolates, but not in any hospital isolates. The ST1 MRSA isolate from the pig sample carried copies of *aac6-Aph2* as well as *ermB*, which was located on a plasmid (Figure 5B). A part of this plasmid which included the *aac6-Aph2* and *ermB* genes was identical to part of GI-PF1, suggesting a shared origin. The shared region (region 2, Figure 5A) contains *ermB* and *aac6-Aph2*, different kinds of transposases such as the Tn3 family transposase and transposase IS1216 as well as the putative transposon Tn552 DNA-invertase bin3. One side of the shared region (region 1) contains genes of a tetracycline efflux MFS transporter (*tetL*), a plasmid recombination/mobilization protein (*pre/mob*) and a plasmid replication protein (*repU*). The remaining part of the island, region 3, has the cadmium resistance *cadDX* operon and the *cst* operon functioning as hydrogen sulfide detoxification. Besides the IS1182 family transposase genes, DNA-invertase gene *hin* and Tn552 DNA-invertase gene *bin3* were also detected in region 3. GI-PF1 region 1 could be detected in 98% (312/318) of *tetL* positive isolates. The distribution of GI-PF1 in the ST398 phylogeny is included in Figure 1.

The HA-VFs *sak, sprD* and *scn* as well as the virulence gene *chp* (chemotaxis inhibitory protein), were located in a prophage region designated phiH1, which has a high level of identity with the *φ3* phage, *Staphylococcus* phage 23MRA (KJ452292.1, 99.43% identity, 89% coverage). The phiH1 prophage could be detected in 54% (144/268) of hospital isolates, but not in pig farm isolates other than the five ST1281 isolates (CC20, all MSSA, one from a farm worker and four from pigs on the same farm).

The MRSA ST398 isolates from pig farms and hospitals had different SCC*mec* elements which are labelled PF-SCC and H-SCC respectively are illustrated in Figure 6. The H-SCC had high similarity with the associated portions of the type V-VT SCC*mec* of the CA-MRSA ST59 JCSC7190 isolate from Chinese Taiwan (98.89% identity) and the type Vc SCC*mec* of the LA-MRSA ST398 reference strain S0385 (98.83% identity). In comparison to the previously reported similar SCC*mec* elements, the PF-SCC appeared to have resulted from a homologous recombination event between the type-IX SCC*mec* of a ST398-t034 MRSA JCSC6943 (Genbank: AB505628.1) from Thailand, and a ST398-t034 MRSA RIVM3897 (Genbank: CP013621.1) from the Netherlands. PF-SCC was detected in all three ST398 MRSA isolated from pig farms. The J1 region of PF-SCC had the genes for the type I restriction and modification system, *hsdR, hsdS*, and *hsdM*, while its J2 region had the *cadDX* operon, *arsRBC* operon and the copper-transporting ATPase gene *copB*.

**Figure 6.**
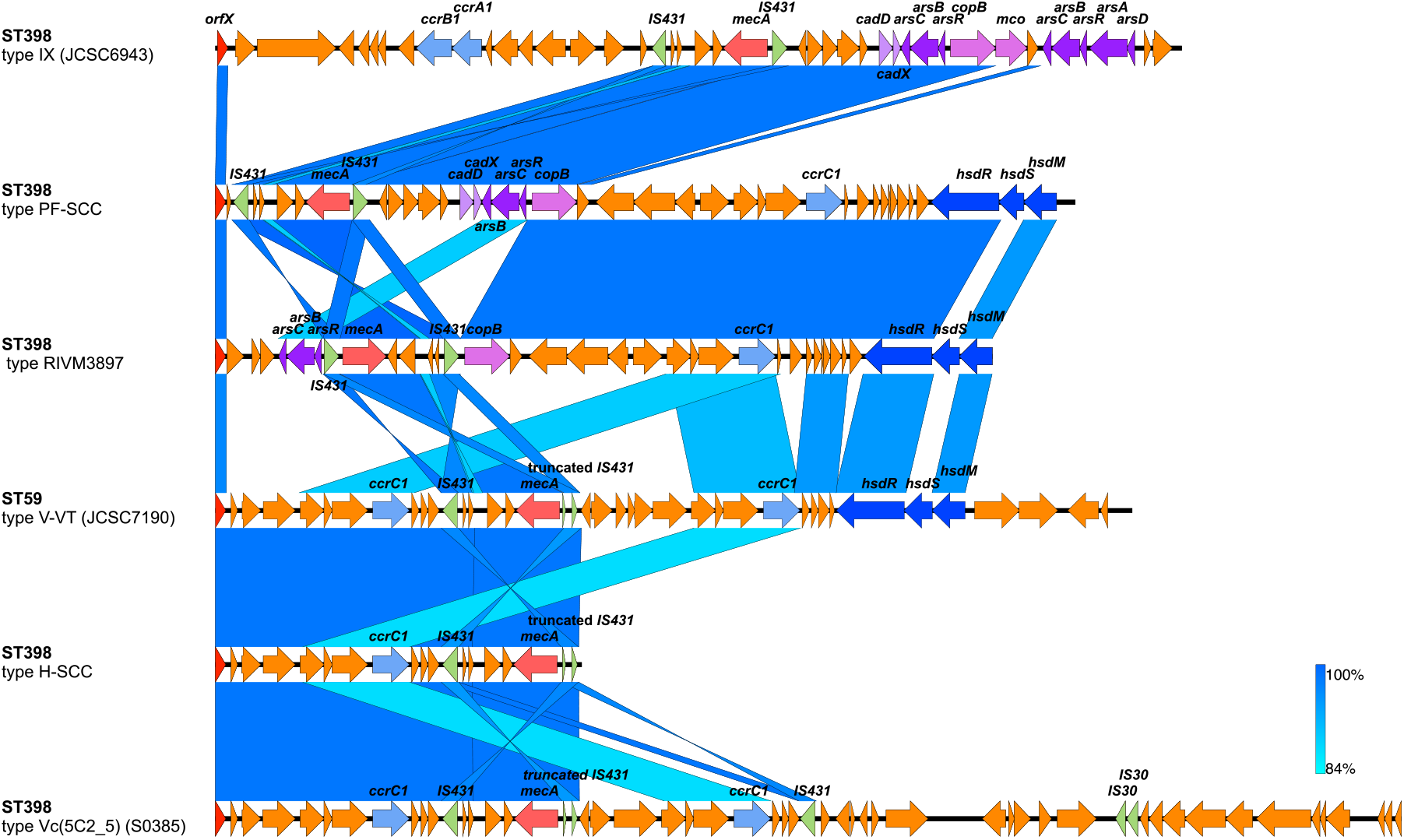
Structures of SCC*mec* elements identified in ST398 MRSA isolates. The diagram shows the main genetic elements and regions of similarity between the SCCmec regions found in MRSA in this study with reference SCCmec types. Sources of the SCCmecs were the hospital MRSA ST398 isolate JL28 (H-SCC), and the pig farm MRSA ST398 isolate 0213-M-4A (PF-SCC). Type V SCC*mec* from ST59 strain JCSC7190 (GenBank: AB512767), type IX SCC*mec* from ST398 strain JCSC6943 (GenBank: AB505628), the SCC*mec* from MRSA ST398 strain RIVM3897 (GenBank: CP013621.1; the SCC*mec* element was determined from 33795 to 69446 bp) and SCC*mec* from ST398 strain S0835 (GenBank:AM990992.1; the SCC*mec* element was determined from 33806 to 88218 bp) are shown for comparison. Regions of similarity (85% to 100%) between each pair of elements are linked with blue bars where the intensity of shading illustrating the degree of similarity.

## Discussion

The striking finding from this study was the discovery that MRSA were present in such a high proportion of pig farms while at the same time there was no evidence of shared populations of MRSA in human and animal hosts. This provides evidence for the potential for these farms to be a source of antibiotic resistant *S. aureus* for human infection in China as is the case for LA-MRSA in Europe but no evidence that LA-MRSA are being found in the human healthcare system in China. In Europe, other than a few countries with low MRSA like Norway and Ireland, LA-MRSA are well established in pig farms with farm level prevalence rates between 25.4% and 100.0% ^29, 30, 31^. These rates are comparable with the MRSA prevalence found in this study although they result from ST398 rather than CC9. Lower rates of CC9 MRSA have been seen in previous studies of Chinese pig farms and slaughterhouse workers^18,32^. While ST398 was found on 3 Chinese pig farms, it was only well established on a single farm as a MSSA lineage. All the ST398 isolates from this study belong to a lineage distinct to the LA-MRSA lineage prevalent in the Europe. Interestingly, this ST398 lineage has been previously been found in China (Figure 1) ^19^ but doesn’t appear to have become established. In comparison, CC9 was the dominant *S. aureus* lineage and appears well established in the majority of pig farms from this study, suggesting that CC9 has a competitive advantage over ST398 in the Chinese pig farm environment. CC9 MSSA isolates are not uncommon in European livestock with small numbers of CC9 MRSA reported^29^, suggesting that the reverse is true there. Further investigations into those differences may provide useful insights into the factors driving host species adaptation which enable LA-MRSA to become established in pig herds.

It is not surprising that out of 42 farm workers carrying MRSA all but 4 carried CC9 MRSA. As is seen in ST398 LA-MRSA, this carriage is likely to be transient, but it does demonstrate how these CC9 MRSA can easily spread from farms to the wider human population. It was therefore surprising not to find any CC9 MRSA among the MRSA isolates collected from the 4 collaborating hospitals in the same region as the farms. It should be noted that it is possible that a larger, more rigorous, or more systematic sampling framework might have found more hospital isolates that might have provided evidence of overlapping populations, in other words, ‘absence of evidence’ is not the same as the ‘evidence of absence’.

Both MRSA and MSSA ST398 were found in the farms and among the hospital isolates which on the basis of MLST suggested that there may have been movement of this lineage between the animal and human populations. The phylogeny of the ST398 isolates in this study together with the wider collection of ST398 *S. aureus* genomes revealed that the hospital ST398 MRSA (C-HM) comprised a separate population to those found on the pig farms (C-L) (Figures 1, and 2). The clade C-HM is widely distributed throughout China including isolates from Central, Eastern and Southern regions of the country. The pig farm ST398 MRSA isolates fall within the diversity of the clade C-L consisting mainly of other livestock isolates from China, predominantly MSSA. The livestock ST398 MSSA came from a single pig farm and were also part of this clade. The hospital ST398 MSSA fall in the diversity of another separate clade. The conclusion from this phylogenetic study is that the animal and human populations of ST398 *S. aureus* in China belong to distinct lineages and there is no evidence that either of the MRSA lineages found on farms is contributing to human MRSA carriage or infection.

The time-scaled phylogeny of an international collection of ST398 isolates (Figure 2) provides a global context revealing the likely evolution of this sequence type. Previous studies had indicated that the prevalent LA-MRSA ST398 lineage had arisen from a human adapted MSSA, become livestock adapted, acquired SCC*mec* and became established on livestock farms, particularly in pig herds^8^. This study shows that the prevalent LA-MRSA shares a common ancestor with a Chinese human MSSA clade (Clade C-H3-2), and that these clades share a common ancestor with the human MSSA clade (Clade C-H3-1) that gave rise to the ST398 MRSA clade (Clade C-HM) from which the hospital isolates from this study came. This topology revealed the diversity of ST398 lineage in Chinese human community and farm animals, highlighting the concern of the emergence of new prevalent CC398 lineages. Given that East Asia was the part of the world where the extant ancestors of the domesticated pig came from, it is possible to speculate that these animals may have been involved in the evolution of this bacterial species.

The investigation of ARGs and VFs identified genes that characterised the MRSA found on farms and in human hospitals. As other studies have found, the human immune evasion cluster of genes found on the *φ3* phage appear to associate strongly with the source of the isolate being human (238/268 hospital isolates carried at least one of the genes). Similarly, the ARGs *tetL, fexA* and *aac6-Aph2* were strongly associated with a farm origin (322/335 pig farm isolates carried at least one of the genes). Part of this may be a simple consequence of clonal expansion from different founder populations but the identification of a genomic resistance island region containing *aac6-Aph2* and *ermB* in most farm isolates which is identical to a region of a plasmid carried by a ST1 MRSA isolate from a pig shows how readily ARGs can move within *S. aureus* populations. The consistency and persistence of both the PF-ARGs and the HA-VFs also indicate a continued selection pressure for the retention of these genes. Examination of the SCC*mec* elements also revealed some consistent differences between the human and pig farm ST398 MRSAs. The SCC found in the hospital isolates (the H-SCC in Figure 6) was a truncated version of a variant of a Type V SCC*mec*. The source could have been type V-VT (sequence from a ST59 CA-MRSA isolated from Chinese Taiwan) or type Vc (5C2&5) (sequence from an ST398 LA-MRSA isolated from the Netherlands). The truncation results in a length of the SCC*mec* which is typical of SCC*mec* types that are associated with CA-MRSA^21,33^. The SCC*mec* found in the farm ST398 MRSA may have resulted from a recombination of ST398 associated SCC*mec* type IX and one found in the isolate RIVM3897 (a human ST398 isolate from the Netherlands). These distinctive SCC*mec* types may provide useful population markers as they currently seem to be restricted to two Chinese clades at present.

In conclusion, the presence of such high levels of MRSA isolates on pig farms is of considerable concern. The most likely driver of this is the overuse of antimicrobials in livestock which has been identified as an issue in China ^34,35^. Fortunately, this study found no evidence that MRSA from pig farms were contributing to the burden of human MRSA found in samples submitted to clinical microbiology laboratories and while ST398 MRSA were found in farms and hospitals there was no shared population that would indicate that transmission had occurred.

## Methods

### Sample collection

Fifty-five pig farms within a 200 km radius of Wuhan City in central China were visited during 2017-2018. For each pig farm, nasal swabs of sows, fattening pigs and farm workers, as well as dust samples were collected. If there were sufficient pigs available on each pig farm, at least 6 sows and 20 fattening pigs were randomly selected and sampled (all the pigs on the farm were sampled if numbers were lower than this). All consenting farm workers were sampled anonymously. Nasal swabs were collected using a transport medium (COPAN Diagnostics Inc.). Environmental dust samples were collected using the following method. Three or four sterile tubes containing 10 ml sterilized Mueller Hinton broth (Oxoid, Ltd.) supplemented with 6.5 % NaCl were opened and exposed in the environment of each selected pig house for at least 10 mins before resealed.

Two city hospitals and two county hospitals in the same region were included in this study. The collaborating hospitals retained all clinical MRSA isolates (for the county hospital H3, MSSA isolates were also retained) that had been identified in the course of normal clinical diagnostics during the same period as the pig farm sampling. The sampling of animals and people complied with protocols subject to ethical review by Huazhong Agricultural University (for pig sampling: HZAUSW-2016-013; for human sampling: HZAUHU-2016-006).

### *S. aureus* isolation and identification

All nasal swabs and dust samples were cultured in Mueller Hinton broth with 6.5 % NaCl, at 37°C for 16 h with shaking. For each culture, a full loop of the culture was spread onto MRSA selective plate (CHROMagar Chromogenic Media) and incubated at 35°C for 24 h. Putative MRSA colonies on the selective plate appeared to be pink coloured. For each sample, up to 3 pink colonies were selected for further isolation and identification. The MRSA candidate isolates were biochemically profiled using the Phoenix 100 ID/AST System (BD Biosciences); those with positive results were stored as MRSA isolates. All the identified *S. aureus* isolates were further sequenced. The species and AST of clinical *S. aureus* isolates obtained from hospitals were also checked using the Phoenix 100 ID/AST System.

### Genome sequencing, assembly and annotation

Genomic DNA was prepared from all the isolates grown overnight at 37°C in tryptone soy broth (BD Biosciences), using bacterial DNA kits (Omega Bio-Tek). All genomic DNA samples were qualified with an optical density at 260 nm (OD_260nm_)/OD_280nm_ ratio between 1.8 and 2 (NanoDrop; Thermo Scientific). Genomic DNA (typically 500 ng) was used to prepare multiplexed libraries for sequencing on Illumina HiSeq 2000 instruments operated according to the manufacturer’s instructions with 100 cycle paired-end runs. For all the sequenced genomes, Fastq_screen^36^ was used to map the raw reads to the most common laboratory contaminants to check for contamination as well as published genomes of *S. aureus* to confirm that all isolates are identified as *S. aureus*. Genome assemblies of all the isolates were generated using SPAdes 3.11.1^37^. Assemblies with an N50 less than 15,000 were discarded. For assembled data, CheckM (version 1.0.13)^38^ was used to analyse the contamination of assembled genomes, the genomes with higher than 1.0% contaminated and 0% strain heterogeneity were excluded. For the six isolates selected for complete genome sequencing, the hospital isolates JL42, JL28 as well as the pig farm isolates 0213-M-4A, 0610-H-2A and 0316-H-5A were further sequenced with Nanopore instruments, while the hospital isolate S82 was sequenced with Pacbio instruments. The sequencing procedures were carried out following the manufacturer’s instructions.

Unicycler(0.4.8)^39^ was used for complete genome assembly, the Illumina sequencing data were used as short reads files and the Nanopore (Pacbio) sequencing data were used as long read files. For the parameters used in Unicycler, the Bridging mode chosen was Normal, which means moderate contig size and mis-assembly rate, and the expected linear sequence number was set to 0.

For all assembled genomes, Prokka (version 1.12)^40^ was used for genome annotation.

### Antimicrobial resistance genes identification

The antimicrobial resistance gene (ARG) database ARG-ANNOT (version 4.0) ^41^ was used for ARGs screening. BLAST+^42^ was used for mapping with the annotated genomes for the identification of ARGs. Sequences with a homology of 90% and an alignment length of 80% of the corresponding reference gene were considered alleles.

### Virulence factors and host-specific genes identification

A database of 216 reported virulence factors (VFs), including virulence, host-specific and other associated genes was created using published research (Table S3). For the sequences mapped to the genes in the database with BLAST+, homology of 90% and an alignment length of 80% were used as cut off value for identification.

### Determination of the pig farm and hospital associated VFs or ARGs

All the *S. aureus* isolates, both MRSA and MSSA, were included in the analysis. Based on the distribution of the VFs or ARGs in the pig farm and hospital *S. aureus* isolates, if an ARG or VF was detected in more than 70% clonal lineages of the pig farm collection and in less than 30% clonal lineages of the hospital collection, as well as in more than 70% isolates of pig farm collection and in less than 30% isolates of hospital collection, the ARG was named as pig farm associated ARG (PFA-ARG) or pig farm associated VF (PFA-VF), conversely, it was named as hospital associated ARG (HA-ARG) or hospital associated VF (HA-VF).

### Multilocus sequence typing

A database including all published alleles of the seven housekeeping gene fragments for *S. aureus* multilocus sequence typing (MLST) downloaded from [https://pubmlst.org/] was built to run the BLAST+ process. BLAST results yielding 100% homology were treated as the same alleles.

### Isolates selected for phylogenetic analysis

A total of 10,535 assembled *S. aureus* isolates were downloaded from the NCBI genome database (date up to Sep. 2019), MLST analysis was used to identify the clonal lineage of the isolates, all ST398 (for Chinese isolates, CC398 were all included) isolates were selected. 77 Chinese CC398 isolates from a previously published^21^ were additionally included. The data was downloaded from the NCBI SRA database, and was then assembled using SPAdes 3.11.1^37^. Combined with the ST398 isolates from this study, a final collection consisted of 904 CC398 isolates. For all these CC398 isolates, CheckM was used to evaluate the assembly genomes, the genomes with higher than 1.0% contaminated and 0% strain heterogeneity (222 isolates) were excluded. According to the BioSample record of each ST398 isolate, 100 isolates with missing location or host information were also excluded. All 115 Chinese CC398 isolates (this study = 36, NCBI genome database = 37, NCBI SRA database = 42) were used for Chinese ST398 phylogenetic analysis. The isolates without the collection date record were also excluded, leaving 582 ST398 isolates (this study = 35, NCBI genome database = 505, NCBI SRA database = 42) as the international ST398 collection (Table S2).

### Phylogenetic analysis of the Chinese CC398 isolates

Snippy (version 4.4.5) [https://github.com/tseemann/snippy] was used to do the genome alignment of the 110 Chinese ST398 isolates. The pig-farm ST398 MRSA in this study 0213-M-4A was used as the reference. Then, for the alignment file, a recombination-removal tool Gubbins (version 2.3.5)^43^ was used to predict the recombination regions. All the recombination regions were marked as “N”, and for the alignment results, all the sites with more than one “N” or gap were trimmed off. For the filtered alignment result, RAxML-NG (version 0.9.0)^44^ was used to build the phylogenetic tree with 500 times bootstrap. A MRSA ST36 strain^8^ (GenBank: BX571856.1) as well as a MRSA ST45 strain (GenBank: CP006044.1) were used as outgroups respectively to double check the root of the tree.

### BEAST analysis of international ST398 isolates

To determine the international phylogeny and molecular clock of ST398 isolates, BEAST 1.8.2^22^ was used to perform the calculation. For the international ST398 collection (n = 582), a RAxML-NG tree was built following the pipeline as described above. Based on the phylogenetic tree, all the clades whose average branch length distance was below 0.00005 were collapsed. Each of the collapsed, as well as the un-collapsed, clades was treated as an independent lineage. For the isolate(s) in each lineage, if the collected date and host and location information was the same, only one of the isolates was selected. Thus, 208 isolates (this study = 11, NCBI genome database = 168, NCBI SRA database = 29) were selected for the analysis with BEAST. Temporal signal of the 208 isolates was investigated using R script provided in the Murray et al., (2015)^45^. The results showed the positive correlation between genetic divergence and sampling time. For the alignment file of the 208 isolates (0213-M-4A was the reference), the recombination regions were predicted with Gubbins and then masked as “N”, and all the sites containing “N” were removed. To account for different evolutionary processes acting at synonymous, non-synonymous, RNA, and non-coding sites^46^, the evolutionary model was partitioned into first and second, third, non-coding and RNA sites according to the reference strain 0213-M-4A. SNPs for each group and the number of the invariant nucleotide sites in each group were calculated, and the information was included in the analysis. For all partitions, we used an HKY + Γ substitution model with the uncorrelated lognormal relaxed clock model.

### Phylogenetic analysis of the Chinese ST9 isolates

For clarity, the isolate number from the study was reduced - for isolates from the same pig farm and same host, that have the same ARGs and VFs patterns and with less than 30 SNPs, only one isolate was selected. The CC9 isolate 0213-P-3B with complete genomic sequence was used as the reference genome and a MRSA ST5 strain (GenBank: NC_002745.2) as well as a ST97 isolate NCTC10344 (GenBank: LS483324.1) (not show in figure) were used as outgroups respectively to root the tree. The other approaches were the same as the phylogenetic analysis of the Chinese CC398 isolates that mentioned above.

### The analysis of canSNP

The presence of three canonical SNPs (canSNP) was identified for each ST398 genome as described by Stegger et al., (2013) ^10^ to determine if isolates were members of the human or livestock clades of LA-MRSA.

### *SCCmec* identification

The reference sequences of each reported *SCCmec* elements from I to XIV were collected as a database for *SCCmec* identification (Table S4). The mapping region with the identity higher than 80% in different contigs of a genome sequence were accumulated to calculate the SCC*mec* type coverage. For each isolate, from all the mapped reference *SCCmec* elements with higher than 70% overall coverage, the one with the highest coverage was identified as the *SCCmec* type of the isolate. The *SCCmec* typing results were double checked with the SCCmecFinder ^47^, the results of the two methods were consistent.

### Identification of other genomic elements

Other genomic elements of the six ST398 isolates with complete genomic sequence were analysed to look for prophages and islands. PHASTER ^48^ was used to predict any prophage region, while IslandViewer 4 ^49^ was used to detect any genomic island region. Searches for genomic elements including prophages, genomic islands, plasmids, and transposons among the whole *S. aureus* collection in this study was performed using BLAST to map regions of interest to the genome sequence of each isolate using an 80% threshold for identity.

### Accession numbers

All the *S. aureus* genomes sequenced in this study are available under the NCBI BioProject: PRJNA660925.

## Acknowledgements

This work was funded by the National Natural Science Foundation of China (NSFC) grant 81661138003 (to R. Z.), and the UK Medical Research Council grant MR/P007201/1 (to M.A.H., A.J.G. and A. W. T.).

